# Visual experience instructs dendrite orientation but is not required for asymmetric wiring of the retinal direction selective circuit

**DOI:** 10.1101/2019.12.19.873364

**Authors:** Malak El-Quessny, Kayla Maanum, Marla B. Feller

## Abstract

Changes in dendritic morphology in response to activity have long been thought to be a critical component of how neural circuits develop to properly encode sensory information. Here we report the impact of dark-rearing on the dendritic morphology and function of a retinal ganglion cell type, a ventral-preferring direction-selective ganglion cell (vDSGC). vDSGCs have asymmetric dendrites oriented along their preferred direction. We found that, at eye opening, vDSGC dendrites are not yet ventrally oriented, and that, surprisingly, dark-rearing prevents ventral orientation of vDSGC dendrites. Despite their dramatic change in dendritic morphology, vDSGCs in dark-reared mice maintain ventral directional preference. Direction selective tuning in dark-reared mice is mediated by asymmetric inhibition, as observed in vDSGCs of normally reared animals. Hence, we postulate that dendritic form follows proper circuit function, where dendritic orientation is refined over the course of development and is dependent on structured visual experience following eye opening.

## INTRODUCTION

Neural computations rely upon precise wiring, which emerges during development. A classic example of such a computation is direction selectivity (DS). In the retina, direction selective ganglion cells (DSGCs) fire many action potentials in response to stimuli moving in a preferred direction (PD) and few to no action potentials in response to stimuli moving in the opposite, or null direction (ND) (Barlow & Levick, 1965). The DS computation is based primarily on the asymmetric wiring of an inhibitory interneuron, the starburst amacrine cell (SAC), onto DSGCs such that motion in the DSGC’s null direction generates more inhibition than motion in the preferred direction (*for review see* Demb, 2007; Vaney and Taylor, 2002). This asymmetric wiring is present as early as postnatal day 10 (P10), a few days prior to eye opening (Wei et al., 2011; Yonehara et al., 2011) and is a consequence of DSGCs preferentially forming synapses with SACs located on their null side, relative to their preferred side (Morrie and Feller, 2015).

What instructs this asymmetric wiring? The establishment of wiring specificity in the nervous system is a complex process, which likely involves the coordination of multiple elements: molecular cues driving synaptic specificity, the geometry of pre- and postsynaptic processes and activity-dependent refinement. In the retinal DS circuit, the relative role of presynaptic (SAC) or postsynaptic (DSGC) cells for instructing this wiring remains a mystery.

A few studies implicate SACs in instructing asymmetric wiring. SACs have radially symmetric processes and serial EM reconstructions show that the orientation of individual SAC process is tightly correlated with the null direction of the DSGCs that synapse onto them (Bae et al., 2018; Briggman et al., 2011; Ding et al., 2016). SAC-specific genetic deletion of the cell-adhesion protein Protocadherin G (Pcdhg) (Kostadinov and Sanes, 2015; Lefebvre et al., 2012) or the axon guidance protein Semaphorin 6A (Sema6A)(Sun et al., 2013), both of which alter SAC radial morphology, eliminate directional tuning of DSGCs. However, in the case of Sema6A, loss of DS is due to a reduction in asymmetric inhibition while asymmetric wiring is maintained (Morrie and Feller, 2018). Interestingly, a hypo-morphic mutation in the FRMD7 gene, which is associated with congenital nystagmus in humans, abolishes direction selectivity along the horizontal axis without impacting SAC morphology (Yonehara et al., 2016). The mechanism by which FRMD7 or other molecules expressed by SACs instruct the selective wiring to different subtypes of DSGCs remains unknown.

An alternative hypothesis is that the postsynaptic DSGCs are instructing asymmetric wiring. DSGCs that encode motion in different directions have distinct molecular profiles and morphological characteristics (Kay et al., 2011; Trenholm et al., 2011). Across the nervous system, the shape of a dendrite has implications for the organization of synaptic inputs as well as its functional role within a circuit (*for review see* Richards and Hooser, 2018; Wong and Ghosh, 2002). Indeed a recent study indicated that the relative orientation of dendrites and axons were more important than molecular identity in instructing synapse specificity in the sensory-motor circuits in spinal cord (Balaskas et al., 2019).

To address the role of DSGCs in the wiring of DS circuits would be to alter the morphology of the DSGC and assess its consequences on DS tuning. To do this, we used the Hb9-GFP mouse line, which expresses GFP in ON-OFF ventral-preferring DSGCs (vDSGCs). Uniquely among DSGCs, vDSGC’s dendritic fields are asymmetric and oriented in their preferred direction across the surface of the retina (Sabbah et al., 2017; Trenholm et al., 2011), making them more likely to form synapses with SACs on their null side. We found that in mice that were dark-reared from birth to adulthood, dendrites of vDSGCs were not oriented towards the ventral direction. Hence, we have used dark-reared vDSGCs to ask whether altered DSGC dendritic morphology influences the wiring and tuning of the DS circuit.

## RESULTS

### Asymmetric dendrites of ventral preferring DSGCs are not ventrally oriented at eye opening or in dark-reared adults

To characterize the impact of dark-rearing on the dendritic morphology of vDSGCs, we filled GFP+ cells in retinas dissected from adult Hb9-GFP mice that were normally-reared (NR) in a 12 h light:12 h dark cycle in adulthood (>P30) and at the time of eye opening (P13/14) (Figure 1a). Dendritic morphology was characterized by a vector pointing from the soma to the dendritic center of mass (dCOM) whose magnitude (ρ) corresponds to the degree of dendritic asymmetry and whose angle (ϴ) corresponds to dendritic orientation (see Figure S1 and Methods).

**Figure 1:**
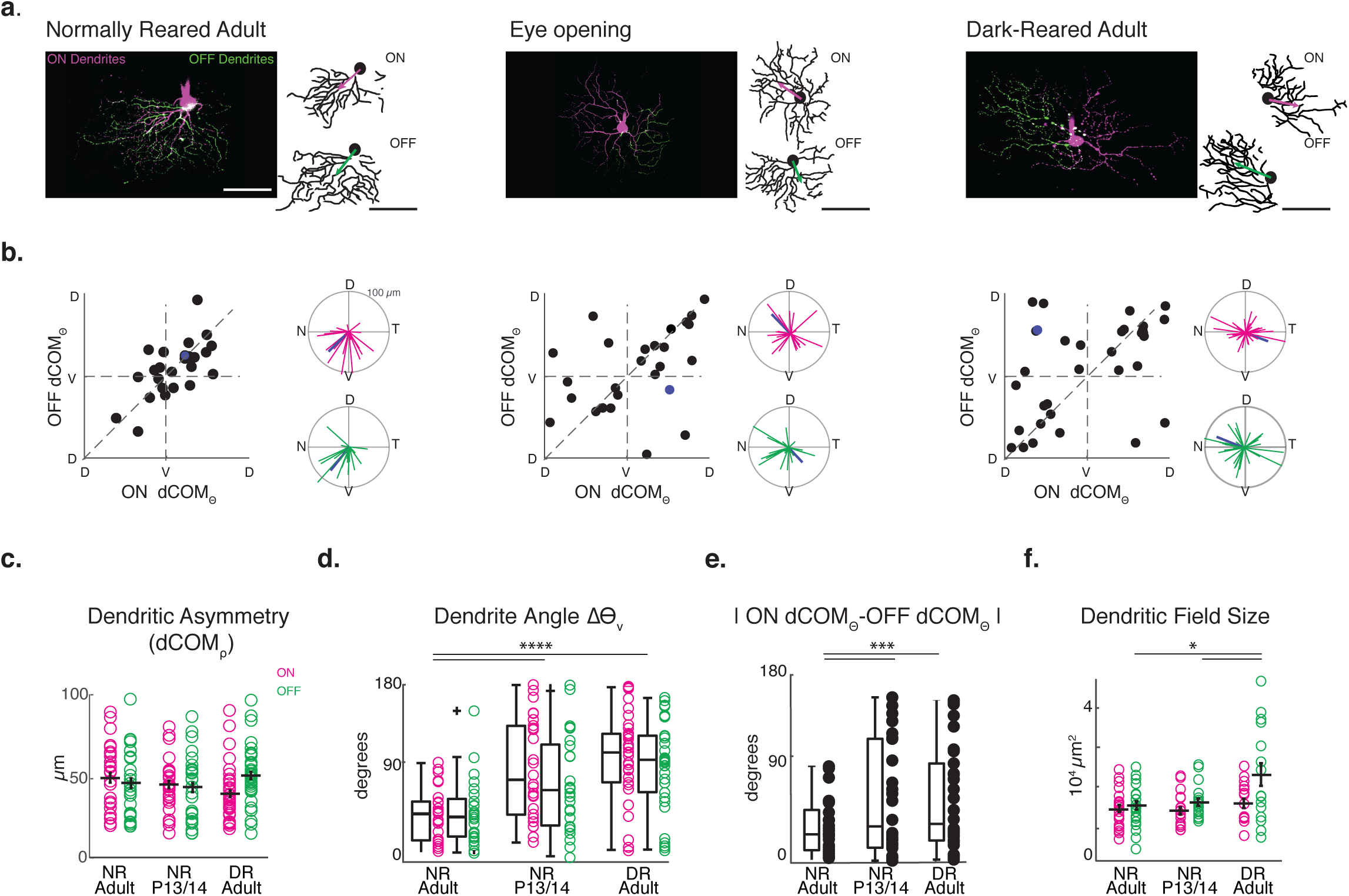
Dark-rearing prevents orientation of vDSGC dendrites toward the ventral direction. **a**. Example maximum intensity projections (left) and binarized skeletons (right) of filled vDSGCs with ON (magenta) and OFF (green) dendrites segments for normally-reared adults (left, P30-50,NR Adult), at eye opening (middle, NR P13/14) and in adults that are dark-reared from birth (right, P30-50, DR Adult) with ON and OFF dCOM vectors overlaied on dendritic skeleton. Scale bar = 100 *µ*m. **b**. Characterization of dendritic orientation in NR adults, at eye opening and in DR adults. Left: ON vs. OFF dendrite center of mass vector angle (dCOM_Ѳ_). Right: polar plots of ON (magenta) and OFF (green) dendrite center of mass vectors. Blue data points refer to example cell above. **c**. Summary data for dCOM vector magnitude as a measurement of ON (magenta) and OFF (green) dendritic asymmetry. Horizontal bar = mean; error bar = SEM. Significance assessed by Kruskal-Wallis one-way ANOVA, p_ON_ and p_OFF_>0.05. **d**. Summary data for deviation of ON (magenta) and OFF (green) dendritic angle from the ventral (270°) axis for all three conditions. Horizontal bar = median. Box plots represent variance. Kruskal-Wallis one-way ANOVA, Dunn-Siddak post-hoc test. ****p<0.0001 **e**. Summary data for absolute difference between ON (magenta) and OFF (green) dendritic angle. Horizontal bar = median. Box plots represent variance. Significance assessed by Levene’s test for absolute variance ***p<0.001 **f**. Summary data for ON and OFF dendritic field size of vDSGCs in three conditions. horizontal bar = mean; error bar = SEM. Significance assessed by one-way ANOVA, p=8.7×10-5; Tukey-Kramer post-hoc test. * p<0.05

As described previously (Trenholm et al., 2011), NR adult vDSGC dendrites were asymmetric, and both ON and OFF dendrites were oriented toward the ventral direction, with ON and OFF dendrites oriented within 45 degrees of each other (Figures 1a and b). However, at P13/14, vDSGC dendrites were asymmetric (Figure 1c), though they were not preferentially oriented towards the ventral direction (Figure 1d). Moreover, the ON and OFF dendrites within a single cell were not always aligned with each other (Figure 1e). In mice that were dark-reared (DR), vDSGCs dendrites retained the eye-opening distribution: asymmetric (Figure 1c) but not preferentially oriented in the ventral direction (Figure 1d) and the ON and OFF dendrites were not always aligned with each other (Figure 1d, Figure S1). Hence, visual experience is necessary for the orientation of both ON and OFF vDSGC dendrites to the ventral direction of the retina.

We also looked at other features of vDSGC dendrites after dark-rearing. We found that dark-rearing increased the dendritic field size of the OFF-stratifying dendrites of vDSGCs (Figure 1f), similar to the impact of dark rearing on OFF-stratifying asymmetric J-RGC (Elias et al., 2018). We also found that the changes in dendritic orientation had no impact on mosaic organization of vDSGCs (Figure S2), indicating that their spacing may not be set by homotypic interactions between dendritic segments (Kay et al., 2012; Rockhill et al., 2000).

### Ventral motion preference is preserved in dark-reared vDSGCs despite altered dendritic morphology

Does function follow form? In NR mice, vDSGC dendrites are oriented in the same direction as their motion preference, where they receive stronger null direction inhibition from SACs located ventral to the DSGC soma (Bae et al., 2018; Briggman et al., 2011; Ding et al., 2016; Morrie and Feller, 2015). Hence, we hypothesized that motion preference would follow dendritic orientation such that vDSGCs with dendrites oriented away from the ventral direction would also have similarly displaced motion preferences. To test this prediction, we conducted cell-attached recordings from vDSGCs and recorded spikes in response to drifting bars moving in 8 different directions in NR adult and P13/14, as well as DR adult animals (Figure 2a). Surprisingly, we observed no differences in the strength or orientation of DS tuning in any of the experimental groups (Figures 2b and 2c). Note, that OFF responses were less tuned at eye opening (Figure 2c), consistent with findings that OFF DS circuits mature later than ON circuits (Hoon et al., 2014; Rosa et al., 2016). Thus, we only analyzed ON spike tuning for P13/14. The preservation of ventral motion preference in P13/14 and adult DR vDSGCs (Figure 2d and Figure S3a) resulted in a strong dissociation of dendritic orientation (dCOM_ϴ_) and spiking preferred direction (PD) (Figure 2d). Moreover, there was no correlation between the size of the dendritic deviation (dCOM_ϴ_) and preferred direction orientation (PD_ϴ_) from the ventral direction (ΔΘ_v_) (Figures 2d, S3a and S3b). Hence, direction selective tuning of vDSGCs appears to be independent of the orientation of the dendrites.

**Figure 2:**
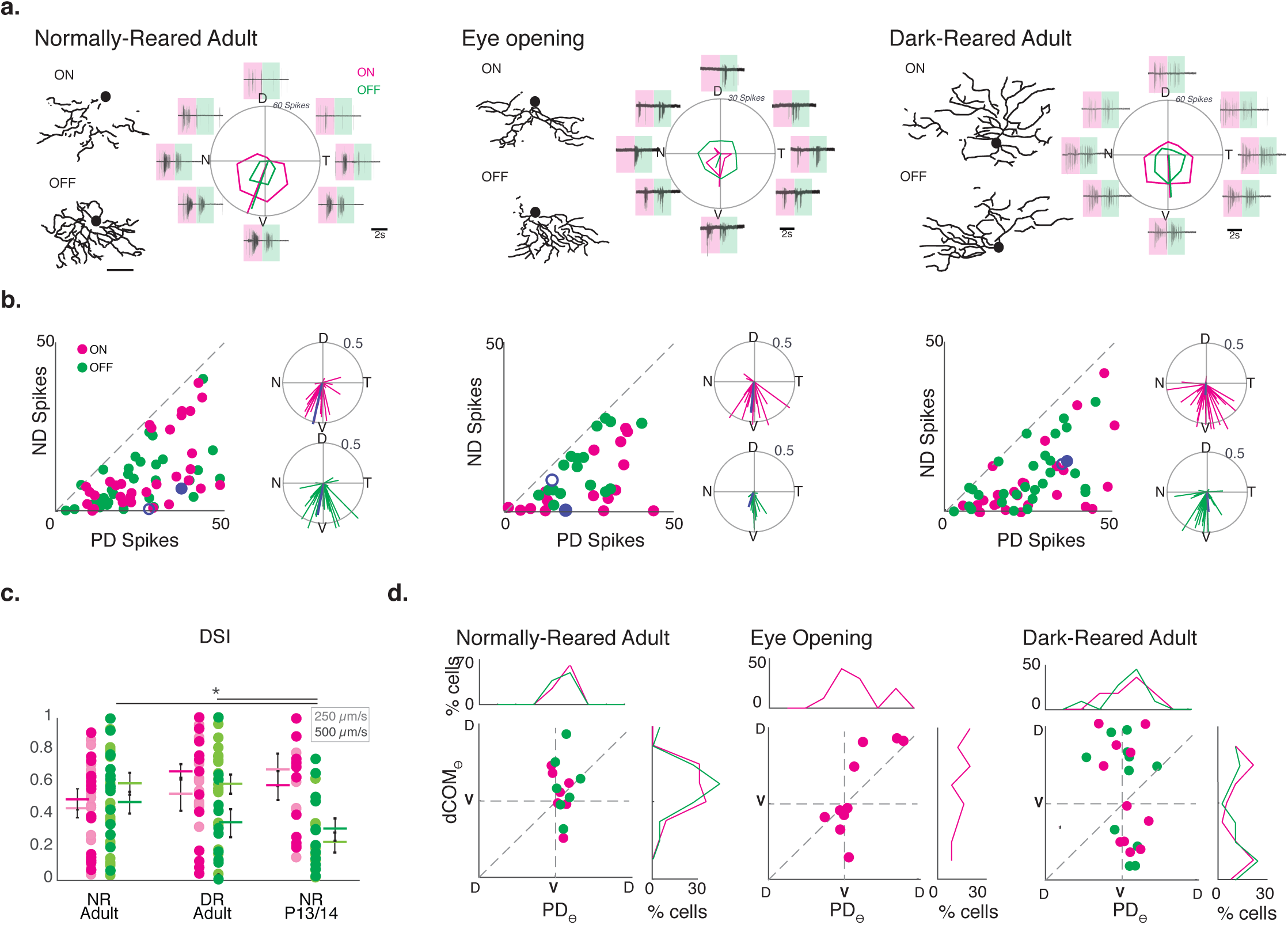
Ventral motion preference is preserved in dark-reared mice despite altered vDSGC dendritic morphology. **a**. Example dendritic skeletons of normally-reared adult (left), eye opening (middle) and dark-reared adult (right) vDSGCs next to the average spike responses of the same cells. Shaded boxes define windows over which ON (magenta) and OFF (green) responses are assessed. Tuning curves plotted in polar coordinates ON (magenta) and OFF (green) responses. Radius of tuning curve = 60 spikes for adult and 30 spikes for eye opening. **b**. Population data represented as number of spikes fired for null direction (ND) vs. preferred direction (PD) spiking (left) and as vectors in polar plots (right inset) of normally-reared adult (left), eye opening (middle) and dark-reared adult (right) vDSGCs. Radius of polar plot: Vector Sum of tuning curve = 0.5, Blue data points refer to ON (filled) and OFF (open) responses of example cell above. **c**. Comparison of spike tuning using direction selectivity index (DSI) at two velocities (lighter shade: 250 *µ*m/s; darker shade: 500 *µ*m/s) for ON (magenta) and OFF (green) responses. Colored horizontal bar =Mean, error bars = SEM. Significance assessed by One-way ANOVA for ON and OFF responses seperately, p_ON_ >0.05, Tukey Kramer post-hoc test * p_OFF_<0.05. **d**. Comparison of dendritic orientation (dCOM_Ѳ_) vs. spiking directional preference (PD_Ѳ_), for normally-reared adult (left), eye opening (middle) and dark-reared adult (right) vDSGCs. X and Y axes centered on ventral axis (V). Top inset: histogram illustrating vDSGC spiking angle. Right inset: histogram illustrating vDSGC dendritic orientation. Note, P13/14 OFF responses were excluded from this analysis due to weak tuning (See Figure 2c). *Mean ± Standard deviation values in Table 2*.

### Asymmetric inhibition from null side starburst cells is preserved in dark-reared vDSGCs despite altered dendritic morphology

Since ventral motion preference is preserved in DR vDSGCs, we assessed the impact of dark-rearing on the directional tuning of both inhibitory (Figure 3) and excitatory (Figure S4) synaptic inputs using whole cell voltage clamp recordings of vDSGCs. Inhibitory synaptic input was highest for dorsal motion (Figures 3a and b) and had similar directional tuning strength (Figure 3c) in vDSGCs from NR adult, P13/14, and DR adult mice. Hence, asymmetric inhibition in response to null direction motion is independent of visual experience and vDSGC dendritic orientation. In NR and DR adults, we observed small EPSC tuning in the preferred direction, which was absent at P13/14 (Figure S4).

**Table 1:** Dendrite orientation.

|  | Adult NR |  | Adult DR |  | Eye opening |  |
| --- | --- | --- | --- | --- | --- | --- |
|  | ON | OFF | ON | OFF | ON | OFF |
| Orientation (dCOM<sub>θ</sub>) | 235.7±105.1 | 258.31±86.5 | 177.3 ±101.3 | 179.9±121.4 | 182.4±116.6 | 201.9 ± 110.2 |
| (ON dCOM<sub>θ</sub> – OFF dCOM<sub>θ</sub>) | 36.5 ±30.56 |  | 57.24 ± 48.15 |  | 56.22 ± 51.7 |  |
| Asymmetry Index (dCOM<sub>ρ</sub>) | 52.9± 29.0 | 48.5 ± 29.2 | 39.4 ±22.9 | 55.0 ±24.5 | 46.5 ± 23.0 | 43.8 ±28.1 |
| Dendrite angle deviation from 270° (Δθ<sub>v</sub>) | 54.1 ± 45.6 | 45.5 ±10.2 | 93.2±47.4 | 81.3 ± 46.1 | 92.5 ± 49.8 | 76.9 ± 48.1 |
|  | n=25 |  | n=35 |  | n=26 |  |

**Table 2:**
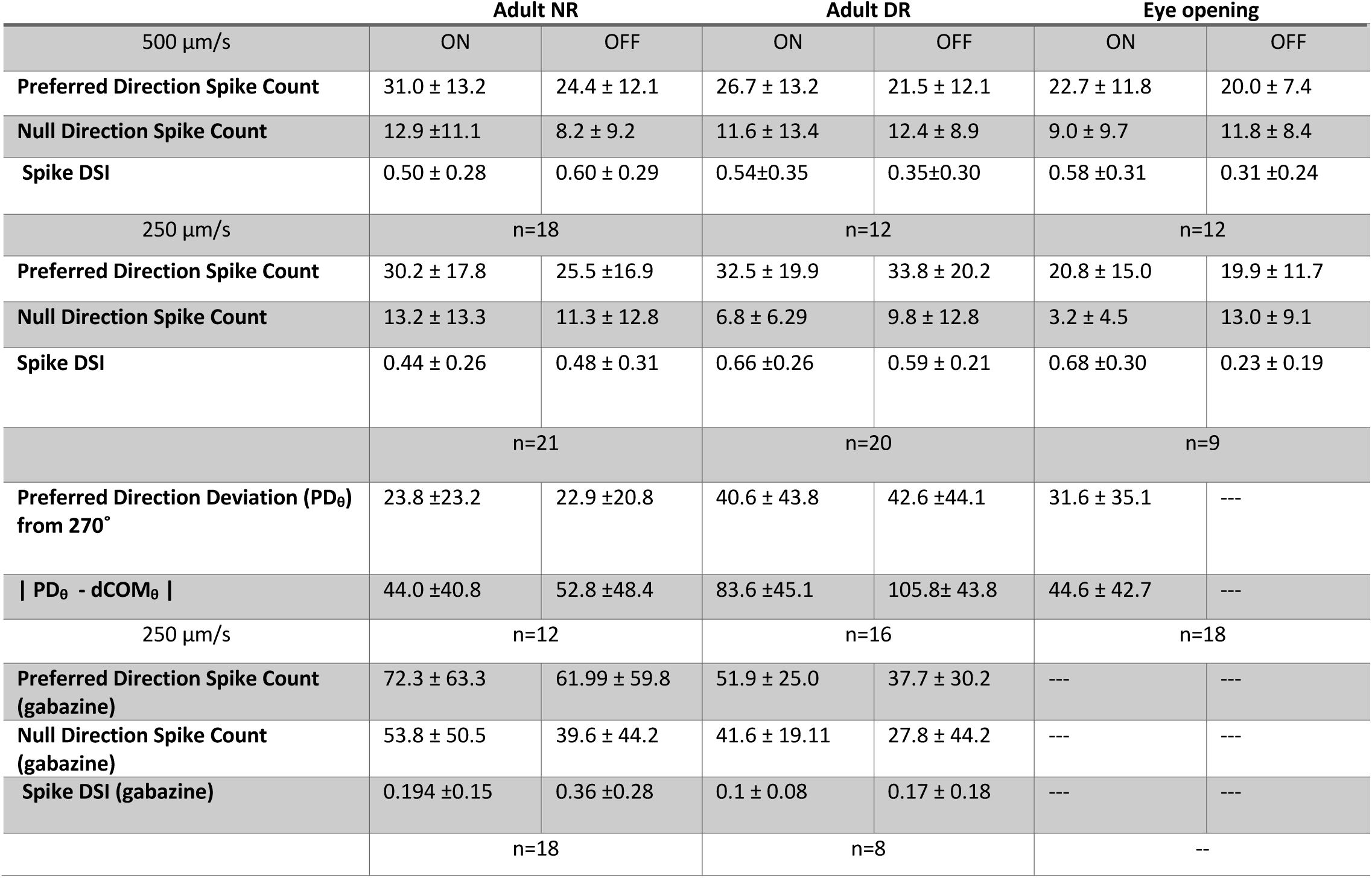
Direction selective tuning.

**Figure 3:**
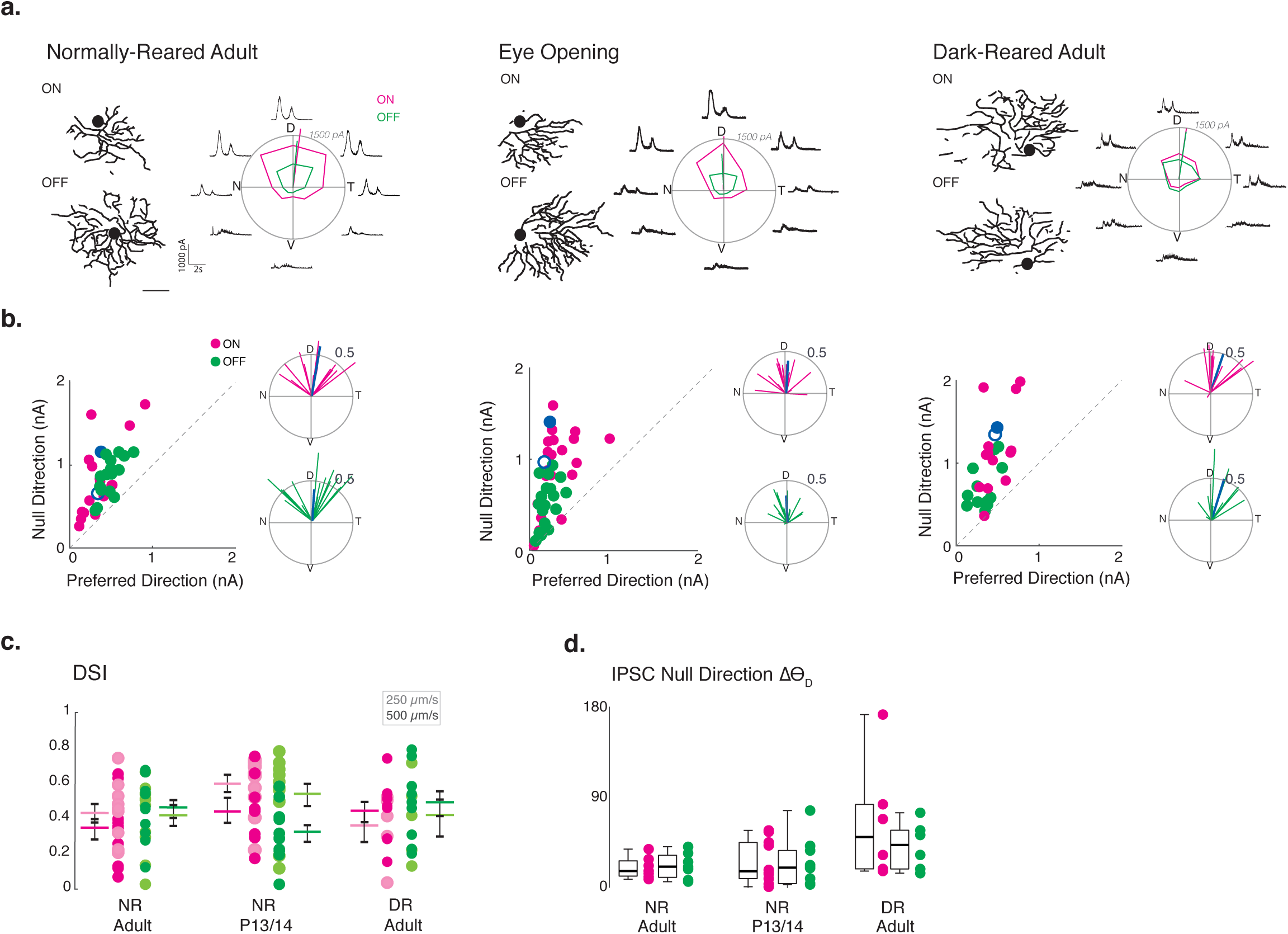
Asymmetric inhibition is maintained in dark-reared vDSGCs. **a**. Example dendritic skeletons of normally-reared adult (left), eye opening (middle) and dark-reared adult (right) vDSGCs next to the average inhibitory currents of the same cells. Tuning curves plotted in polar coordinates ON (magenta) and OFF (green) responses. Radius of tuning curve = 1500 pA. **b**. Left inset: population data for normally-reared adult (left), eye opening (middle) and dark-reared adult (right) vDSGCs represented as peak amplitude of IPSC recorded for null direction (ND) vs. preferred direction (PD) stimulation. Right inset: polar plot for normalized vector sums of population IPSC tuning curves. Radius of polar plot: Vector Sum of tuning curve = 0.5. Blue data points refer to ON (filled) and OFF (open) example cell above. **c**. Tuning strength vDSGC inhibitory input, across two stimulus velocities (lighter shade: 250 *µ*m/s; darker shade: 500 *µ*m/s) quantified as the direction selectivity index (DSI) for normally-reared (NR) adult, eye opening (NR P13/14) and dark-reared (DR) adult. Colored horizontal bar = mean, errors bars = SEM. One-way ANOVA, p>0.05. **d**. Summary data for deviation of preferred direction of IPSC tuning of vDSGCs (ND) from the dorsal (90°) axis for all three conditions. Horizontal bar = median. Box plots represent variance. Kruskal-Wallis test, p>0.05. *Mean ± Standard deviation values in Table 3*.

A feature of the direction selective circuit in the retina is the presence of spatially offset inhibition (Fried et al., 2002; Hanson et al., 2019; Pei et al., 2015; Sivyer et al., 2010). To determine whether directionally tuned IPSCs were generated by SACs located on the null side of vDSGCs, we mapped the excitatory and inhibitory receptive fields of NR and DR adult vDSGCs by recording synaptic currents evoked by squares of light sequentially presented at 100 block-shuffled locations within a soma-centered grid (Figure S5a). We calculated the vector from the soma to the center of the inhibitory and excitatory receptive fields (Figure 4a and b), using the magnitude (ρ) of the vector to indicate receptive field displacement, and the angle of the vector (Θ) to indicate receptive field orientation. We found that in both NR and DR adult vDSGCs, the inhibitory receptive fields were similarly oriented toward the ventral direction relative to the soma (Figure 4c). We then computed the vector from the excitatory to the inhibitory receptive field and observed that the magnitude of their displacement (ρ) from each other was similar across conditions (Figure 4d). Moreover, the angle of the vector was similarly aligned to the ventral axis (ΔΘ_v_) (Figure 4e). Hence, although DR vDSGCs have displaced dendrites, they still receive spatially offset inhibition from SACs located on their null side. These data indicate that the location of dendrites does not influence the selectivity of their presynaptic partners and that antiparallel SAC-DSGC geometry is not necessary for asymmetric wiring of inhibition.

**Table 3.** Synaptic inputs and receptive fields.

| Synaptic Inputs: Figure 3 | Adult NR |  | Adult DR |  | Eye opening |  |
| --- | --- | --- | --- | --- | --- | --- |
| 500 $\mu\text{m/s}$ | ON | OFF | ON | OFF | ON | OFF |
| Preferred Direction IPSC amplitude | 354 ± 228 | 252 ± 128 | 501.3 ± 169.2 | 296.7 ± 148.2 | 348.1 ± 231.3 | 226.6 ± 132.3 |
| Null Direction IPSC amplitude | 875.7 ± 474.8 | 651.7 ± 243.4 | 1279 ± 453.6 | 801.4 ± 261.5 | 861.60 ± 352.1 | 460.4 ± 271.9 |
| IPSC DSI | 0.43 ±0.17 | 0.45 ±0.16 | 0.42 ±0.18 | 0.47 ± 0.22 | 0.45 ±0.24 | 0.31 ± 0.16 |
| 250 $\mu\text{m/s}$ | n=11 | | n=8 | | n=11 | |
| Preferred Direction IPSC amplitude | 451.3 ± 248.5 | 331.7 ± 358.7 | 381.4 ± 38.2 | 267.3 ± 128.5 | 193.10 ± 79.9 | 141.21 ± 56.7 |
| Null Direction IPSC amplitude | 965.2 ± 509.2 | 642.3 ± 320.5 | 862.3 ± 327.9 | 629.1 ± 320.5 | 962.0 ± 469.7 | 573.6 ± 287.6 |
| IPSC DSI | 0.35 ± 0.78 | 0.41 ±0.18 | 0.34 ±0.19 | 0.39 ±0.23 | 0.60 ±0.16 | 0.54 ±0.21 |
|  | n=12 |  | n=5 |  | n=11 |  |
| Null Direction Deviation ($\text{PD}_\theta$) from 90° | 18.8 ± 10.7 | 20.9 ± 12.9 | 6.9 ± 26.9 | 3.5 ± 11.7 | 5.6 ± 14.5 | 5.1 ± 14.3 |
|  | n=7 |  | n=5 |  | n=10 |  |
| Receptive fields: Figure 4 |  |  |  |  |  |  |
| EPSC peak amplitude | 216.9 ± 100.0 | 194.6 ± 79.7 | 199.28 ± 85.83 | 167.44 ± 79.34 | -- | -- |
| IPSC peak amplitude | 484.5 ± 194.1 | 383.4 ± 133.1 | 489.6 ± 143.2 | 367.6 ± 106.3 | -- | -- |
| Inhibition ($\Delta\theta_v$) | 31.9 ± 27.0 | 31.6 ± 29.5 | 34.2 ± 25.2 | 32.9 ± 24.4 | -- | -- |
| E-I distance | 31.6 ± 20.7 | 35.0 ± 31.0 | 53.2 ± 29.3 | 58.6 ± 30.7 | -- | -- |
| E-I Angle | 239.4 ± 76.6 | 247.0 ± 85.6 | 276.9 ± 55.2 | 277.2 ± 28.0 | -- | -- |
| E-I Angle ($\Delta\theta_v$) | 54.5 ± 52.5 | 42.3 ± 33.5 | 47.0 ± 24.9 | 40.0 ± 23.6 | -- | -- |
|  | n=13 |  | n=9 |  | -- |  |

**Figure 4:**
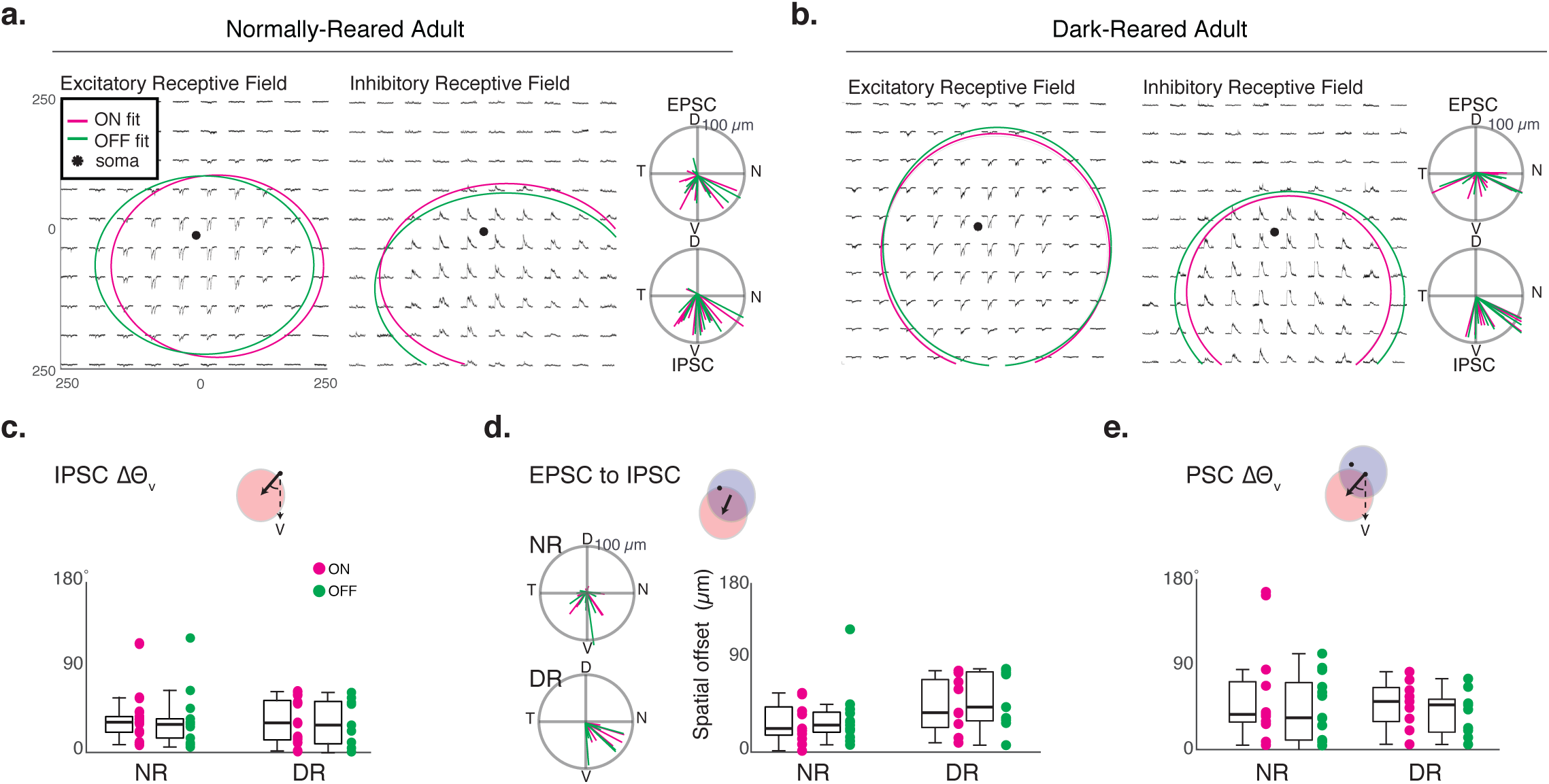
Inhibitory Receptive field is spatially offset from the excitatory receptive field in adult vDSGCs. **a**. Example mean excitatory (left) and inhibitory (right) PSC responses of normally-reared (NR) adult vDSGCs. Peak amplitude of ON (magenta) and OFF (green) responses are fit with a 2-dimensional gaussian. Ellipse radius in X and Y axes = 2 standard deviations of the gaussian fit centered around fit peak. Right: Population data of NR adult vDSGCs represented as polar plots of vectors from the soma to the excitatory (top) and inhibitory (bottom) receptive field fits. Radius of polar plot = 100 *µ*m, center of polar plot = soma location. **b**. Example mean excitatory (left) and inhibitory (right) PSC responses of dark-reared (DR) adult vDSGCs. Peak amplitude of ON (magenta) and OFF (green) responses are fit with a 2-dimensional gaussian. Ellipse radius in X and Y axes = 2 standard deviation of the gaussian fit centered around fit peak. Right: Population data of DR adult vDSGCs represented as polar plots of vectors from the soma to the excitatory (top) and inhibitory (bottom) receptive field fits. Radius of polar plot = 100 *µ*m, center of polar plot = soma location. **c**. Comparison of the orientation of the vector from the soma to the center of the inhibitory receptive field, relative to the ventral axis (ΔΘ_v_). One way ANOVA, p>0.05. **d**. Left: Population data of NR (top) and DR (bottom) adult vDSGCs represented as polar plots of vectors from the excitatory to the inhibitory receptive field centers. Radius of polar plot = 100 *µ*m, center of polar plot = EPSC receptive field center. Right: Comparison of the spatial offset (the magnitude of the vector) from the center ON (magenta) and OFF (green) excitatory receptive field to the center of the ON and OFF inhibitory receptive field fits (indicated in schematic above, where excitatory receptive field fit is in blue and inhibitory receptive field fit is in red) in NR and DR adult vDSGCs. One way ANOVA, p >0.05. **e**. Comparison of the orientation of the vector from the center of the excitatory receptive field to the center of the inhibitory receptive field, relative to the ventral axis (ΔΘ_v_). One way ANOVA, p>0.05. *Mean ± Standard deviation values in Table 3*.

### Dark-rearing prevents the maturation of inhibition-independent directional tuning in vDSGCs

The asymmetric morphology of vDSGCs, along with the alignment of their dendritic arbor and preferred direction, has been postulated to underlie their ability to retain some directional tuning in the absence of inhibitory input (Trenholm et al., 2011). Modeling experiments suggest that this postsynaptic mechanism of directional tuning is mediated by nonlinear conductances at the distal dendrites of vDSGCs, which allow them to integrate excitatory input along the soma-to-dendrite ventral direction more efficiently than in the opposite dorsal direction.

To test whether this postsynaptic contribution to DS is maintained with dendrites that are not ventrally oriented, we compared DS tuning in the presence of a GABA_A_ receptor blocker, gabazine (50 µM), in NR and DR mice using stimuli moving at 250 µm/s, which enhances the postsynaptic dendritic contribution to DS in adult NR vDSGCs (Trenholm et al., 2011). In NR vDSGCs, we found that OFF responses retained ∼86% of their DS tuning in the presence of gabazine, while ON responses retained ∼46% of their DS tuning (Figure 5b). The presence of gabazine led to an increase in both preferred and null direction spiking (Figure S5a), which underlie the changes in DSI. In contrast, in DR vDSGCs, DS tuning was nearly absent in the presence of GABA_A_ receptor inhibition, with ON and OFF responses retaining only 16% and 30% of their DS tuning, respectively (Figure 5d). Here, we observe a greater increase in null direction spiking, relative to preferred direction spiking in gabazine (Figure S5b), which underlies the more dramatic decrease in DSI. These observations provide strong evidence that the establishment of the postsynaptic contribution to DS in vDSGCs requires visual experience.

**Figure 5:**
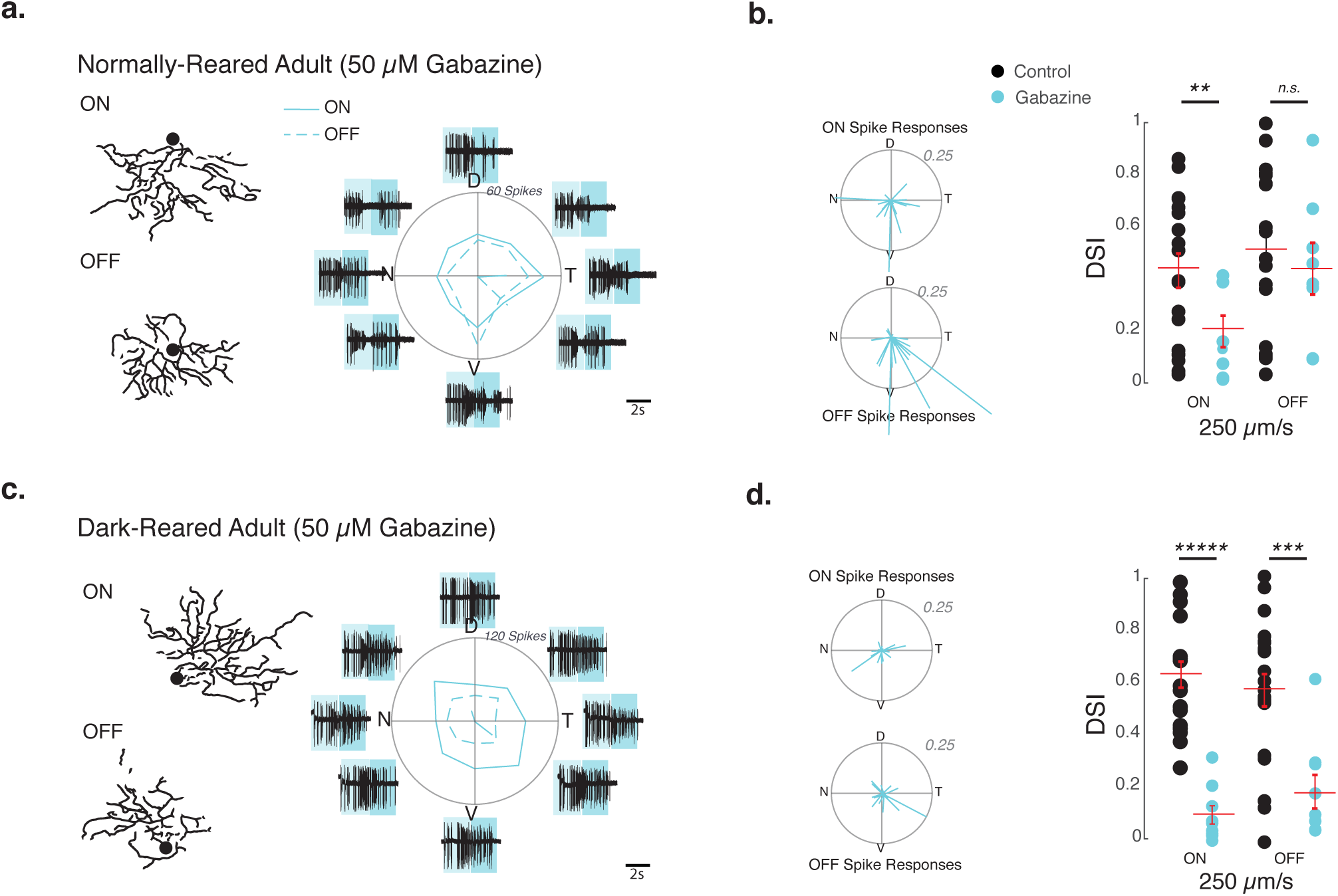
Dark-reared vDSGCs with misoriented dendrites lose directional tuning in the absence of inhibitory input. **a**. Left: example dendrite skeletons for ON and OFF dendritic segments for normally-reared adult vDSGCs. Right: example average spike responses of the same cell in the presence of 50 *µ*M gabazine. Gray boxes define windows over which ON (light blue) and OFF (darker blue) responses are assessed. Tuning curves plotted in polar ON (solid line) and OFF (dashed line) responses. Radius of tuning curve = 60 spikes. **b**. Left: Population data of normally-reared adult vDSGC responses in 50 *µ*M gabazine represented as vectors in polar plots. Data for ON (top) and OFF (bottom) plotted separately. Radius of polar plot = 0.25. Right: Comparison of tuning strength as quantified by the direction selectivity index (DSI) of normally reared adult vDSGCs across using a slow velocity stimulus (250 *µ*m/s) in control conditions (black) and in 50 *µ*M gabazine (cyan). Statistical significance assessed acrosss velocities by one-way ANOVA, p_ON_, p_OFF_ <0.05; Tukey Kramer post-hoc test p-values indicated in figure. Note control data is shared from figure 2c. ** p<0.01 **c**. Left: example dendrite skeletons for ON and OFF dendritic segments for dark-reared adult vDSGCs. Right: example average spike responses of the same cell in the presence of 50 *µ*M gabazine. Gray boxes define windows over which ON (light gray) and OFF (darker gray) responses are assessed. Tuning curves plotted in polar ON (solid line) and OFF (dashed line) responses. Radius of tuning curve = 120 spikes. **d**. Left: Population data of dark-reared adult vDSGC responses in 50 *µ*M gabazine represented as vectors in polar plots. Data for ON (top) and OFF (bottom) plotted separately. Radius of polar plot: Vector sum of the polar plot = 0.25. Right: Comparison of tuning strength as quantified by the direction selectivity index (DSI) of normally reared adult vDSGCs in control conditions (black) and in 50 *µ*M gabazine (cyan). Statistical significance assessed by one-way ANOVA, p_ON_, p_OFF_ <0.05; Tukey Kramer post-hoc test p-values indicated on figure. Note control data is shared from figure 2. *** p<0.001, *****p<10^−5^ *Mean ± Standard Deviation values in Table 2*.

## DISCUSSION

Dendritic morphology is thought to play a critical role in neural computations and is shaped by many factors during development. Here we report that after eye opening, asymmetric vDSGCs orient their dendrites ventrally and that this orientation requires visual experience, as it is prevented by dark-rearing. Dark-rearing selectively disrupted the postsynaptic contribution to DS tuning in vDSGCs but spared the tuning of excitatory and inhibitory synaptic inputs, and, as a result, dark-reared vDSGCs retained their normal directional tuning. This study demonstrates that the precise asymmetric wiring of synaptic circuits that mediate direction selectivity is independent of the orientation of the DSGC dendrite. Hence, the maturation of dendritic morphology appears to be dictated by the functional circuit rather than the traditional view that dendritic morphology dictates circuit function.

### Activity influences dendritic morphology

We found that dark-rearing had a dramatic but specific effect on the dendrites of vDSGCs – it prevented the orientation of vDSGC dendrites in the ventral direction and, in a subset of vDSGCs, prevented the alignment of ON with OFF dendrites (Figure 1). This finding indicates that visual experience provides an instructive cue for vDSGC dendrites to orient themselves along their preferred direction. This counters our initial hypothesis that the orientation of DSGC dendrites along their preferred direction optimizes antiparallel wiring with null side SAC dendrites. Rather, we postulate that the orientation of the dendrites along their preferred direction is instead necessary for the establishment of postsynaptic mechanisms for DS. Here we consider these findings in the context of other systems where afferent activity plays a role in asymmetric dendritic development.

How might visual experience influence vDSGC dendrite orientation? One clue comes from looking at other examples of neurons whose asymmetric dendritic arbors are oriented towards their presynaptic partners, including ganglion cells in the zebrafish retina (Choi et al., 2010), layer IV Stellate cells in the somatosensory cortex (Greenough and Chang, 1988; Nakazawa et al., 2018; Woolsey and Van der Loos, 1970), mitral cells of the rodent olfactory bulb (Blanchart et al., 2006; Hinds and Ruffett, 1973), among others (*for review see* Wong and Ghosh, 2002). In developing zebrafish, retinal ganglion cells (RGCs) initially have both apical and basal dendrites pointing both toward and away from the inner plexiform layer. Upon contact with bipolar cells, RGCs basal dendrites reorient toward the apical side to form functional synapses (Choi et al., 2010). In the *heart-and-soul (has)* mutant zebrafish, RGCs are displaced to the other side of bipolar cells, leading to an increase on basal orientation. This indicates that the location of the RGC dendrites relative to bipolar cell afferents instructs their orientation. In layer IV spiny stellate neurons in barrel cortex, activity-dependent pruning eliminates inactive dendrites oriented away from barrel centers while sparing active dendrites oriented toward the barrel center, which elaborate and stabilize (Harris and Woolsey, 1981; Li et al., 2013; Nakazawa et al., 2018; Narboux-Nême et al., 2012). When thalamocortical axon input is silenced via infraorbital nerve cut, or via postsynaptic NMDA receptor knockout, spiny stellate neuron dendrites misalign with respect to barrel centers (Mizuno et al., 2014, 2018). This suggests that dendrites are oriented toward afferent activity.

Sensory experience may be required for the establishment of dendritic orientation through mechanisms that influence gene transcription. In situ hybridization revealed that the transcription factor BTBD3 is highly localized to the barrels of somatosensory cortex during development and that activity-dependent nuclear translocation of BTBD3 is required for the orientation of stellate neuron dendrites towards the barrel hollows (Matsui et al., 2013). Similarly, BTBD3 is implicated in the orientation of dendrites in ferret visual cortex towards the center of ocular dominance columns whereby monocular enucleation and BTBD3 shRNA knock-down lead to misorientation of layer IV excitatory neuron dendrites (Matsui et al., 2013).

How does visual experience instruct vDSGCs to orient towards their preferred direction? One possibility is that bipolar cell inputs on the preferred side are stronger and therefore dendrites on the preferred side are stabilized while dendrites on the non-preferred side are pruned. Such asymmetric wiring of bipolar cells was recently reported in ON DSGCs (Matsumoto et al., 2019). However, serial EM reconstructions of symmetric ON-OFF DSGCs (Ding et al., 2016) do not indicate such a bias, though this reconstruction has not been conducted for vDSGCs.

Our finding that dark-rearing did not impact stratification differs from previous studies where visual deprivation prevented the stratification of some ganglion cell dendrites (Tian and Copenhagen, 2001) via a process dependent on Brain Derived Neurotrophic Factor (BDNF) activation of TrkC receptors (Liu et al., 2007). Interestingly, this activity dependent stratification does not appear to be dictated by glutamate release from bipolar cells, since vDSGC ON dendrite stratification has been shown to rely on the expression of the adhesion molecule Contactin 5 and its co-receptor Caspr4 for proper stratification, which is expressed by ON SACs (Peng et al., 2017), and since stratification was normal in mice lacking release from ON bipolar cells (Kerschensteiner et al., 2009).

### Implications for postsynaptic contributions to DS

Thus far, ventrally preferring asymmetric DSGCs (vDSGCs) have been the only DSGC subtype to exhibit directional tuning in the absence of inhibitory input (Trenholm et al., 2011). In the presence of the GABA_A_ receptor antagonist gabazine, other subtypes of anterior and posterior preferring DSGCs lose their directional tuning (Ackert et al., 2009; Bos et al., 2016; Rivlin-Etzion et al., 2011; Wei et al., 2011). However, due to the correlation of dendritic orientation and preferred direction in normally-reared adult vDSGCs, it is postulated that they also possess postsynaptic mechanisms for encoding motion direction. Our results in normally-reared adults are consistent with previous findings, where vDSGCs retain around 50% of their tuning in the absence of inhibitory input (Trenholm et al., 2011). By using a bar stimulus in our experiments, we were able to discriminate ON and OFF responses more precisely than the moving spot stimulus used in previous experiments. We found that 86 % of OFF tuning is retained while only 46% of ON tuning is retained, indicating the postsynaptic contribution to direction selectivity in vDSGCs is stronger in the OFF dendrite.

Computational modeling experiments have shown that vDSGCs may possess nonlinear conductances, like voltage gated sodium channels, on their distal dendrites allowing them to efficiently encode ventral motion (from soma to distal dendrites) more efficiently than dorsal motion (from dendrites to soma) (Trenholm et al., 2011). More recent work has shown that gap junction coupling between dendrites of neighboring vDSGCs promotes dendritic spikes in post-junctional cells, in response to synchronous synaptic input (Trenholm et al., 2013; Yao et al., 2018) suggesting that gap junctions may serve as a nonlinear conductance underlying the postsynaptic DS. Whether visual experience is critical for the maturation of the source of nonlinear postsynaptic conductance, e.g. voltage gated conductances or electrical coupling, or the organization and kinetics of excitatory inputs onto DSGC dendrites (Matsumoto et al., 2019) remains to be determined.

## METHODS

### Animals

Mice used in this study were aged from p13-60 and were of both sexes. To target ventral preferring DSGCs, we used Hb9::GFP (Arber et al., 1999) mice, which express GFP in a subset of DSGCs (Trenholm et al., 2011). Normally-reared animals were kept on a 12h:12h dark-light cycle. Dark-reared animals were kept on a 24h:0h dark-light cycle from birth until tissue collection. All experiments involved recording from 1-7 cells from at least 3 animals of either sex. All animal procedures were approved by the UC Berkeley Institutional Animal Care and Use Committee and conformed to the NIH Guide for the Care and Use of Laboratory Animals, the Public Health Service Policy, and the SfN Policy on the Use of Animals in Neuroscience Research.

### Retina Preparation

Mice were anesthetized with isoflurane and decapitated. Retinas were dissected from enucleated eyes in oxygenated (95% O2/5% CO2) Ames’ media (Sigma) for light responses or ACSF (in mM, 119 NaCl, 2.5 KCl, 1.3 MgCl2, 1 K2HPO4, 26.2 NaHCO3, 11D-glucose, and 2.5 CaCl2) for paired recordings. Retinal orientation was determined as described previously(Wei et al., 2010). Isolated whole retinas were micro-cut at the dorsal and ventral halves to allow flattening, with dorsal and ventral mounted over two 1–2 mm2 hole in nitrocellulose filter paper (Millipore) with the photoreceptor layer side down, and stored in oxygenated Ames’ media or ACSF until use (maximum 10 h). All experiments were performed on retinas in which dorsal-ventral orientation was tracked.

### Retinal Histology

Whole-mount retinas were fixed in 4% PFA for 20 min, then washed in block solution (2% donkey serum, 2%bovine serum albumin, 0.3% Triton X-100 in PBS, 3 times, 16 min). Next, retinas were incubated in primary antibodies (1:1000 rabbit anti-GFP, Invitrogen, Grand Island, NY; 1:500 goat anti-ChAT, Millipore, Billerica, MA) for 1-3 days, and then washed in block solution (3 times, 15 min) and left in block solution at 4°C overnight. The retinas were then incubated in secondary antibody (1:1000 donkey anti-rabbit Alexa Fluor 488, 1:1000 donkey anti-goat Alexa Fluor 568; Invitrogen) at 4°C overnight. Then, they were washed in block solution (5 times, 30 min) and left in PBS overnight. Then, retinas were mounted and cover slipped with Vectashield (Vector Laboratories, Burlingame, CA).

### Two-photon microscopy and morphological reconstruction

After physiological recordings of vDSGCs were completed, Alexa-594-filled vDSGCs in the Hb9– GFP mice were imaged using the two-photon microscope at 700nm. At this wavelength, GFP is not efficiently excited but Alexa 594 is brightly fluorescent. 600 × 600 µm Image stacks were acquired at z intervals of 1.0 um and resampled fifteen times for each stack using a 20X objective (Olympus LUMPlanFl/IR 2x digital zoom, 1.0 NA) 30kHz resonance scanning mirrors covering the entire dendritic fields of the vDSGCs. Image stacks vDSGCs were then imported to FIJI (NIH) and a custom macro was used to segment ON and OFF dendrites based on their lamination depth in the inner plexiform layer (ON layer 10-30 um, OFF layer 35-65 um depth). Following ON and OFF dendritic segmentation, another custom FIJI macro uses the local maximum values of fluorescent pixels to binarize and skeletonize ON and OFF dendritic segments for morphological analyses.

### Dendrite morphological analysis

To assess morphological alignment of vDSGC dendrites in oriented retinas, we calculated the center of mass of the dendritic pixels from the binarized vDSGC skeleton relative to the soma. Briefly, a FIJI macro was designed to do the following: 1) allow user to localize the soma, record soma coordinates, and then clear/exclude soma pixels from the stack and dendritic analysis 2) creates a maximum intensity projection of the dendritic pixels and measure their center of mass (COM) using the following equation:

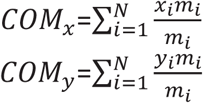

where N is the total number of dendritic pixels, *x* is the coordinate distance of each pixel and *m* is the mass of each pixel *i*, which is either 1 or 0.

3) Following COM calculation, the length of the vector from the dendritic COM to the soma coordinates is used as a measurement of dendritic asymmetry, and the angle of the vector from the dendritic COM to the soma coordinates is used as a measurement of dendritic angle.

Note: Dendritic reconstructions shown in figures were obtained by manually tracing example cells using the simple neurite tracer plugin on FIJI. Dendritic skeletons were then rendered and eroded for presentation.

### Whole retina morphological analysis

Fixed and stained whole-mounted retinas were imaged on an epifluorescent macroscope (Olympus MV PLAPO 0.63x) within one week of mounting. Exposure and gain were adjusted per retina to maximize GFP signal. Images were then analyzed on FIJI for use in mosaic and nearest neighbor analysis (Figure S2). Whole retina fluorescent images were processed through a custom built MATLAB script. Briefly, somas were selected and a mask of the retinal outline was defined. Then, the nearest neighbor distance (NND) was calculated for every soma, and the distance was divided by the total size of the retinal outline to normalize for age-dependent size differences. The regularity index (or conformity ratio) was calculated by dividing the mean NND by the standard deviation from the mean (Riemann, 1978). Both NND and regularity index metrics were compared to a random distribution of the same number of somas for each retina.

For dendritic area, the analyzer was blind to age and rearing condition of each cell. Furthest dendritic extent was determined by adjusting brightness and contrast of individual maximum intensity z-projection images. For mosaic analysis, cell bodies of v-DSGCs were manually marked for each retina to create binary masks, and outlines of retinal areas were manually traced. In addition, a random array of points (representing the same number of somas on that retina) within the boundary mask of each retina was generated in MATLAB. We used a custom MATLAB script to calculate nearest neighbor distances of our masked somas and compared these distances to the random distributions. All statistical analyses were performed in MATLAB. Student’s two-tailed t-test was used to compare across rearing groups and across age groups. All ± SEM unless otherwise noted.

### Two-photon targeted loose patch and whole-cell voltage-clamp recordings

Oriented retinas were placed under the microscope in oxygenated Ames’ medium at 32–34°C. Identification and recordings from GFP+ cells were performed as described previously(Wei et al., 2010). In brief, GFP+ cells were identified using a custom-modified two-photon microscope (Fluoview 300; Olympus America) tuned to 920 nm to minimize bleaching of photoreceptors. The inner limiting membrane above the targeted cell was dissected using a glass electrode. Cell attached voltage clamp recordings were performed with a new glass electrode (4-5 MΩ) filled with internal solution containing the following (in mM): 110 CsMeSO4, 2.8 NaCl, 20 HEPES, 4 EGTA, 5 TEA-Cl, 4 Mg-ATP, 0.3 Na_3_GTP, 10 Na_2_Phosphocreatine, QX-Cl (pH = 7.2 with CsOH, osmolarity = 290, ECl^−^ = −60 mV). After cell attached recordings of spikes, whole cell recordings were performed with the same pipette after obtaining a GΩ seal. Holding voltages for measuring excitation and inhibition after correction for the liquid junction potential (−10 mV) were 0 mV and −60 mV, respectively. Signals were acquired using pCLAMP 9 recording software and a Multiclamp 700A amplifier (Molecular Devices), sampled at 10 kHz, and low-pass filtered at 6 kHz.

### Visual Stimulation

For visual stimulation of vDSGCs, broad-band visible light ranging from 470 to 620 nm was generated using an OLED display (SVGA Rev2 OLED-XL; eMagin) displaying custom stimuli created using MATLAB software with the Psychophysics Toolbox. Drifting bars were presented (velocity = 250 and 500 um/s, length =600 μm width =350 μm over a 700 μm radius circular mask) in 8 block shuffled directions, repeated 3 times, with each presentation lasting 6 s and followed by 500 ms of grey screen) were projected through the 20X water-immersion objective (Olympus LUMPlanFl/IR 360/1.0 NA) onto the photoreceptor layer through the same 20x objective used to target cells once the cell attached recording configuration was achieved. The illumination radius on the retina was 1.4 mm to limit modulation of DSGC responses by inhibitory wide-field amacrine cells (Chen et al., 2016).

The directionally selective index (DSI) was calculated for spike responses as: 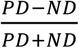 where PD is the number of spikes in the preferred direction and ND is the number of spikes in the null direction. We also used the magnitude of the vector sum of the spike responses as another measurement of directional tuning.

### vDSGC whole cell recordings

For cell attached vDSGC recordings, spike counts were calculated by bandpass filtering traces (0.08-2 kHz) and manually identifying a threshold value for spikes on the filtered traces. Local minima below threshold that did not violate refractory period criteria (0.001 s) were counted as spikes. ON and OFF responses were defined as spikes occurring within a 1.9 s time window starting right before the presentation of the leading or trailing edge of the stimulus. The average spike counts across the 3 trials were used to calculate the vector sum of the spike responses. Preferred directions for both ON and OFF responses used to calculate average spike counts and were defined as the angle of the vector sum of spike responses for the ON and the OFF responses.

For experiments conducted in gabazine (Tocris, SR95531), we diluted 50 µM in AMES media, and allowed it to perfuse for 5-10 mins at a perfusion rate of 1 mL/min.

For voltage clamp recordings, traces were first average across the 3 trials for each direction and inspected to ensure consistency of responses. Average traces were baseline subtracted based on the last 500 ms of recording or a user defined interval after manual inspection. Peak currents were calculated from average baseline subtracted traces and were the maximal (IPSC) or minimal (EPSC) points during the 1.9 s window described above. The peak currents were used to calculate the vector sum of the current responses. Preferred directions for both ON and OFF responses used to calculate peak responses and were defined as 180 ± the angle of the vector sum of ON and OFF peak IPSCs, or the angle of the vector sum of ON and OFF peak EPSCs if IPSCs were not recorded in that cell.

### Receptive field mapping

To map excitatory and inhibitory receptive fields of vDSGCs, visual stimuli were generated using a computer running 420 nm light through a digital micro mirror device (DLI Cel5500) projector with a light emitting diode (LED) light source through a 20X objective (UMPlanFL 0.5NA W). Stimuli (30 μm^2^) at an intensity of 4×10^9^ photons/s/μm^2^ were presented in a pseudorandom order, in a 10×10 grid, onto a stimulus field of 500 μm^2^, with the DSGC soma located in the center of the stimulus field (See Figure 5). Voltage clamp recordings were simultaneously acquired using methods described above.

To quantify excitatory and inhibitory receptive field sizes for each cell, we first divided each trace into the ON and OFF response based on the stimulus we present. Next, we fit a two-dimensional Gaussian to the post synaptic current (PSC) peak values averaged over three trials. We use the 2x standard deviation of the gaussian fit to display the size of the receptive fields. To compare the size and location of the PSC receptive fields relative to the soma and to each other, we used the standard deviation and peak coordinates of the Gaussian fits, respectively.

