## Supplemental Figures for "Visual experience instructs dendrite orientation but is not required for asymmetric wiring of the retinal direction selective circuit"

### Supplemental Figure 1

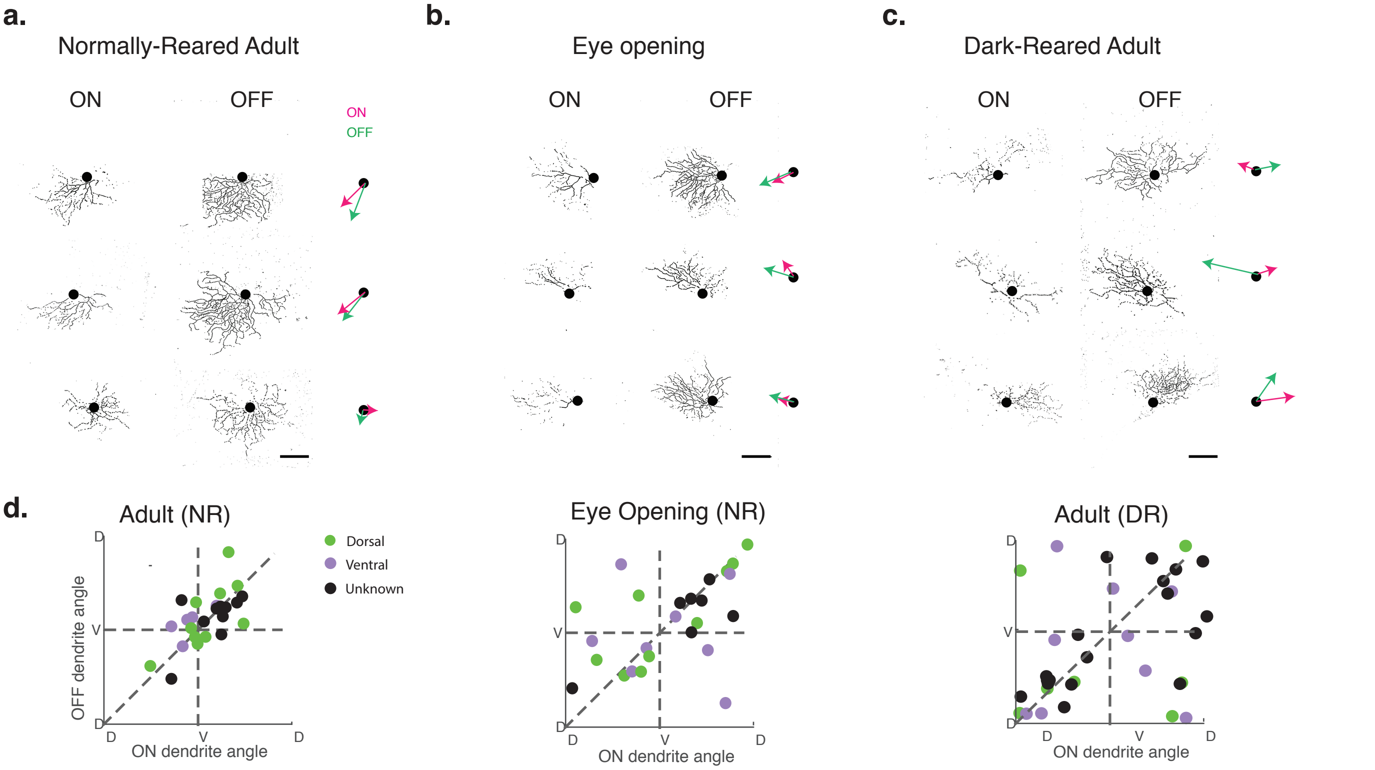


**Figure S1: Dendritic Center of Mass (dCOM) as a measurement for dendrite orientation and asymmetry. (Supplemental to Figure 1)**

**a-c.** Three examples of ON and OFF skeletons of normally-reared adult (a) eye opening (b) and dark-reared adult (c) vDSGCs. Image scale bar = 100 $\mu$m. Right: magenta (ON) and green (OFF) vectors of dCOM about the soma.

**d.** Dendritic orientation in NR adults, at eye opening and in DR adults. Color indicates location on the retina. We do not observe an effect of location on the orientation of ON and OFF dendrites.

### Supplemental Figure 2

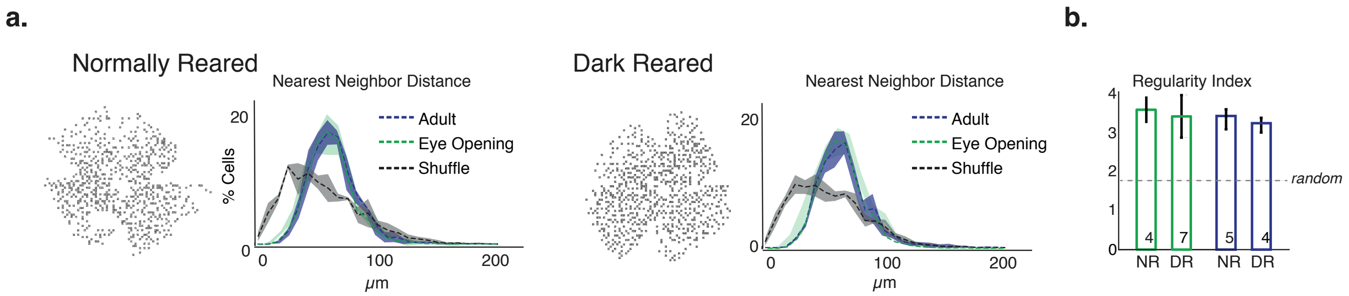


**Figure S2: Visual experience does not alter mosaic organization of vDSGC somas. (Supplemental to Figure 1)**

**a.** Mosaic analysis of soma locations for normally reared vs dark reared adults . Left: binarized image marking soma locations across retina. Right: Nearest neighbor distance distributions (10 $\mu$m bins) for actual locations and for randomly shuffled distances. Significance assessed by one-way ANOVA, $p>0.05$.

**b.** Regularity index shows that vDSGC somas are non-randomly distributed under all conditions. Numbers at the bottom of bar plot = number of retinas sampled. Random regularity index value obtained from a distribution of randomized distances.

### Supplemental Figure 4

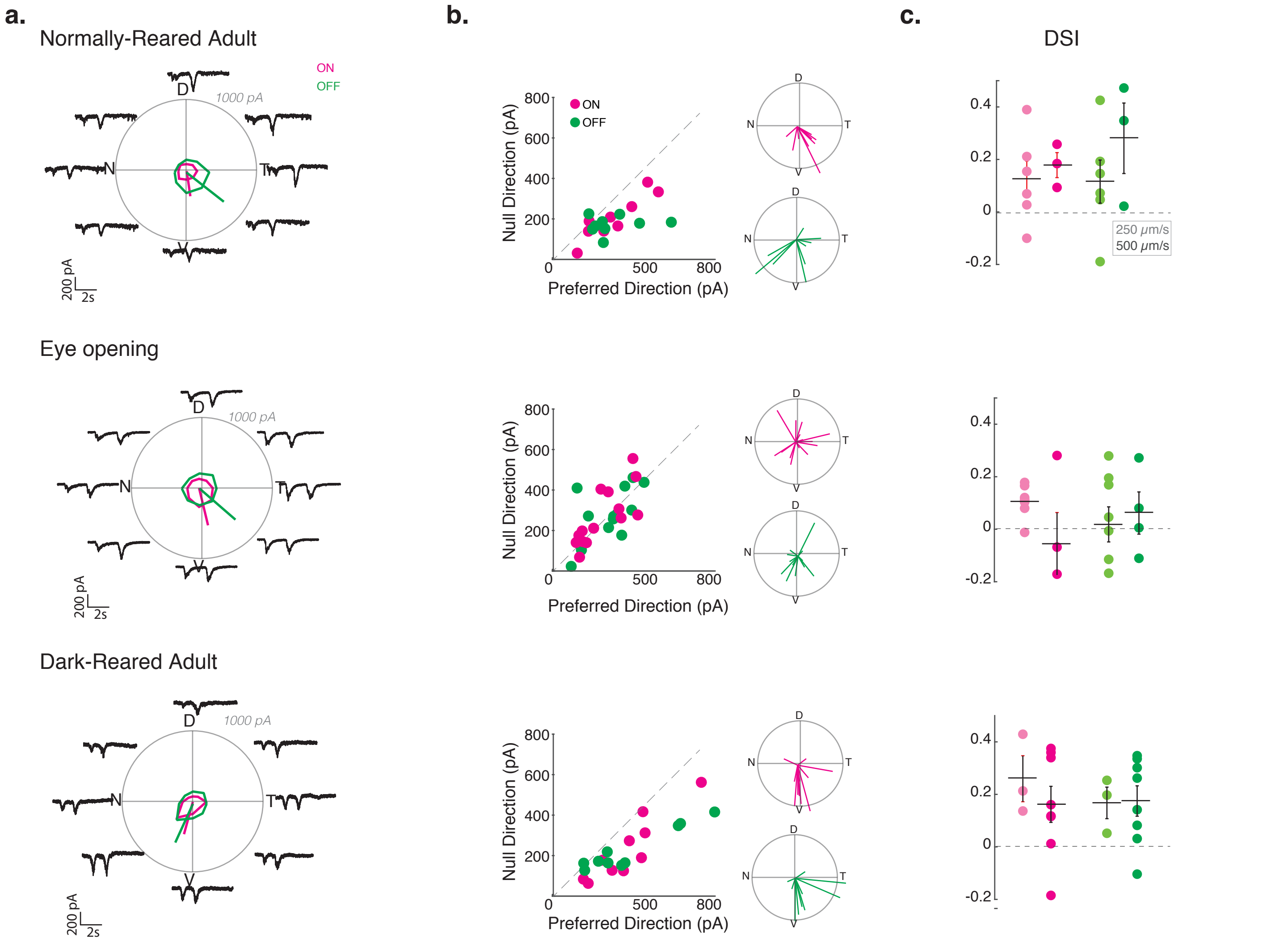


**Figure S4: Adult vDSGCs exhibit weakly tuned excitatory input. (Supplemental to Figure 3)**
**a.** Example tuning curves of average ON (magenta) and OFF (green) excitatory currents of vDSGCs in normally-reared (top), at eye opening (middle row), and in dark-reared adults (bottom row) . Radius of polar plots = 1000 pA.
**b.** Left: population data represented as peak amplitude of EPSC recorded for null direction (ND) vs. preferred direction (PD) stimulation in normally-reared adults (top) eye opening (middle) and dark reared (bottom) adults. Right: polar plot for normalized vector sums of population EPSC tuning curves. Note, radius of polar plot; Vector sum of the tuning curve = 0.25.
**c.** Tuning strength of ON (magenta) and OFF (green) excitatory input of vDSGCs across two stimulus velocities (lighter shade: 250 $\mu$m/s; darker shade: 500 $\mu$m/s) quantified as the direction selectivity index (DSI) Horizontal bar = mean, errors bars = SEM. Significance assessed by t-tests.

### Supplemental Figure 3

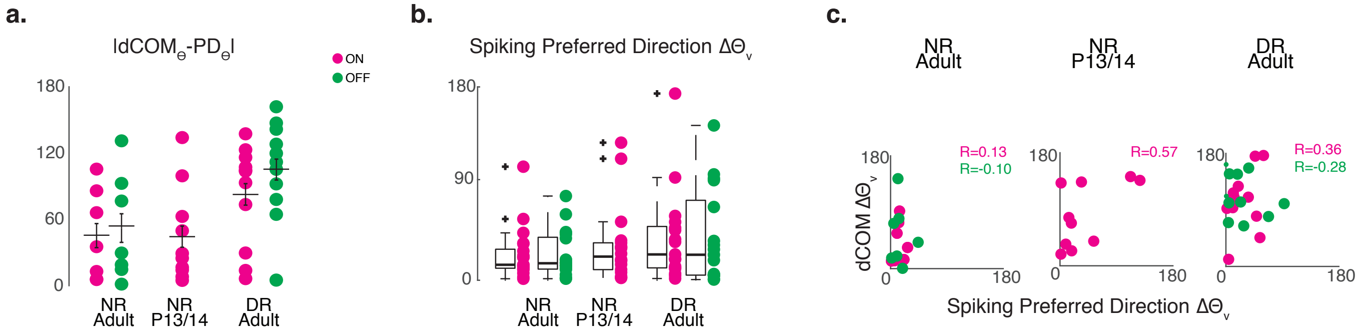


**Figure S3: Ventral motion tuning is preserved in dark-reared mice despite altered vDSGC dendritic morphology. (Supplemental to Figure 2)**

**a.** Quantification of the alignment of dendrite orientation and preferred direction of vDSGCs. Data for ON (magenta) and OFF (green) plotted separately. Horizontal bar = mean, errors bars = SEM. Significance assessed by Kruskal Wallis test, $p>0.05$. Note, P13/14 OFF responses were excluded from this analysis due to weak tuning (See Figure 2c).

**b.** Quantification of the alignment of vDSGC for ON (magenta) and OFF (green) spiking preferred direction to the ventral direction ($\Delta\Theta_v$). Horizontal bar = median. Significance assessed by Kruskal-Wallis test, $p>0.05$. Note, P13/14 OFF responses were excluded from this analysis due to weak tuning.

**c.** Comparison of the Spiking preferred direction and dCOM angle deviation from the ventral axis ($\Delta\Theta_v$). Correlation coefficient (R) for ON (magenta) and OFF (green) values indicated on plot.

### Supplemental figure 5

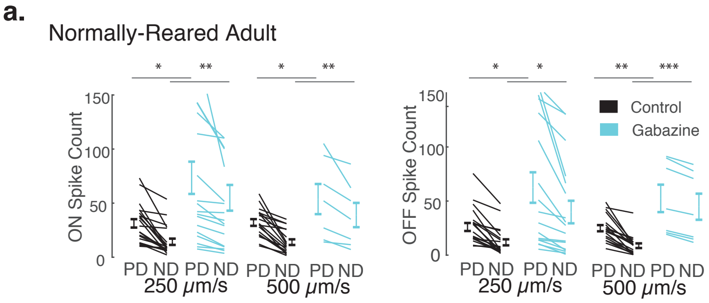


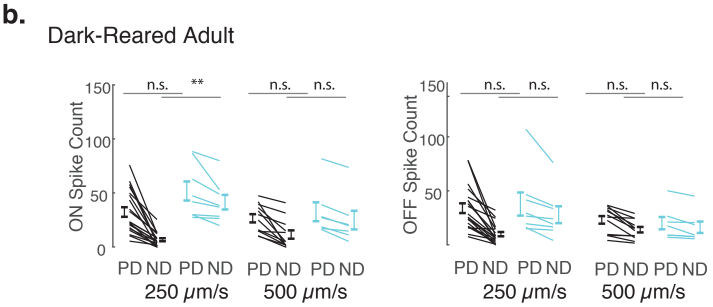


**Figure S5: Normally-reared and dark-reared vDSGCs exhibit greater null direction spiking in gabazine. (Supplemental to Figure 5)**

Preferred (PD) vs. null (ND) direction spike counts for ON and OFF responses of normally-reared (a) and dark-reared (b) adult vDSGCs, across two velocities. Statistical significance assessed by unpaired t-tests. n.s. : $p>0.05$, *$p<0.05$, **$p<0.01$, ***$p<10^{-4}$
